# Unrelated males in colonies of facultatively social bee

**DOI:** 10.1101/2020.01.20.912519

**Authors:** Michael Mikát, Daniel Benda, Jakub Straka

## Abstract

Colonies of social Hymenoptera are usually groups of closely related females, in which the dominant female(s) is specialized for reproduction and subordinate females care for immature offspring. Kin selection is thought to be the main factor that supports social cohesion. We have discovered a simple society of the bee *Ceratina chalybea* with an average of 4.68 colony members that cannot be maintained by kin selection alone. These colonies consisted of old reproductive female, young adults and provisioned brood cells. About half of young adults are unrelated to the old female, and almost all of the young adults are male. The old female provisions new brood cells, while continuing to feed young adult offspring. As young adults do not perform demanding or risky activities, they incur little or no cost, but they do benefit from the food they obtain from the old female.

## Introduction

Cooperation between individuals is one of the most interesting biological phenomena. Several mechanisms of cooperation have been proposed [1–4], but it is thought that kin selection is the main mechanism driving the organization of societies [5–7]. This theory claims that the spreading of alleles through one’s own reproduction is equivalent to the spreading of alleles by related individuals [6,8]. The strength of kin selection is strongly influenced by relatedness – any help provided to another individual is only beneficial if the cost of helping that individual is lower than the benefit of the help received multiplied by the relatedness between the individuals [6,9]. Another mechanism of cooperation that has been postulated is based on reciprocity: the cost of helping is compensated by the predicted future benefit [1,3]. Alternatively, cooperation may be a by-product of selfish behaviour [4]. Recently, the plausibility of group selection has been disputed – sacrificing individuals for group benefit can exist as an adaptive feature [5,10,11].

Colony members in almost all social insects are related [7,9,12]. Cooperation of unrelated individuals occurs under specific conditions and some derived states [13,14]. Pleometrosis (nest founding by multiple females) is one of the most common situations in which unrelated members of a colony cooperate [13,15]. In many wasp and bee societies, some proportion of drifting workers have been recorded [14,16]. These workers originate from foreign colonies and are therefore not related to the original nest members [14]. Unlike eusocial insects, unrelated helpers are common in cooperative breeding vertebrates [2].

Societies of the eusocial Hymenoptera are mostly composed of females because the queens and workers are female [17,18] and most males die soon after mating, not participating in the life of the society [18–20]. Exceptions, such as male participation in care, are rare among eusocial societies and male helpers usually have only a minor role in colony life [21–23].

The traditional division of labour in hymenopteran societies is between the queen, who performs most or all egg-laying, and the workers, who are responsible for other tasks, especially food provisioning [18,24]. However, in small societies that are composed of one reproductively dominant female and one, or a few other females, a different type of task division is possible. The dominant female can perform egg-laying as well as food provisioning, with the reproductively subordinate female(s) performing nest guarding and other tasks in the nest [25]. This system of provisioning by the reproductively dominant female is typical for Xylocopinae bees [25,26].

Here, we examined social nests of the small carpenter bee, *Ceratina chalybea.* Small carpenter bees from the genus *Ceratina* make their nests in broken twigs with soft pith [27]. The female excavates a burrow in the pith and later provisions the brood cells [28,29]. After completion of cell provisioning, the female usually guards the offspring until they reach adulthood [29–31]. Subsequently, she provisions newly emerged offspring with pollen and nectar [31–33]. Although most temperate *Ceratina* are solitary [30,34], they belong to the Xylocopinae family, which is ancestrally facultatively social, with obligate solitary nesting being a derived state [35]. Social nests of *Ceratina* contain a few females, usually two but sometimes as many as four [36,37].

Until now, *C. chalybea* was thought to be a solitary bee [30]; however, here we present the first evidence of social nesting in this species. In contrast to most *Ceratina* bees, the females of this species did not obligately guard their offspring until adulthood, rather females had two alternative strategies – either guarding or abandoning the nest after provisioning of brood cells is finished [30]. Clearly, mothers can only perform provisioning of young adults in guarded nests and only these nests can develop into eusocial colonies. Here, we describe the social nests of *C. chalybea.* Moreover, we tested relatednes between colony members. We try examine possible costs and benefits for reproductive dominant and subordinate colony members.

## Methods

### Study site

We performed field research at the Havranické vřesoviště (coordinates 48°48’32.6"N 15°59’33.6"E) location, near the village of Havraníky, in Podyjí National park. This location is situated in the Southern Moravian region of the Czech Republic. The main experiments were performed in the years 2015 and 2017, but additional data are also presented from the years 2013 and 2018. Office of Podyjí National park permitted this research.

### Preparation of nesting opportunities

We studied nests made in artificial nesting opportunities. The nesting opportunities were made from cut twigs with pith, from the following plant species: *Solidago canadensis*, *Helianthus tuberosus*, or *Echinops sphaerocephalus*. Twenty twigs were tied together into a sheaf and fixed to the ground with a bamboo rod. We distributed more than 20,000 nesting opportunities (1000 sheaves) each year. In 2013 and 2015, we collected nests directly from the nesting opportunities. In 2017 and 2018, some nests were collected directly from sheaves at nesting site and other nests were taken from original sheaves and transport to study plot for observation. Nests were collected and dissected after the observation period.

### Nest dissection

We collected nests in the evening (after 19:00) to ensure that all inhabitants would be present inside the nest. We stored nests in a fridge between collection and dissection. We opened nests using a knife or clippers, and for each nest we recorded the presence of all adults and non-adult juveniles (eggs, larvae and pupae). For adults, we noted sex and age (parental vs filial generation). The age of adults was easily recognized because adults of the parental generation had extensive wear to their wings. All nests had only one old female. For non-adult juveniles, we recorded the stage and position of its brood cell in the nest. We also recorded the number and position of any empty cells (cells without provisions or offspring, [30]). We measured the length of the nest. We distinguished between new nests and reused nests. Reused nests had adult excrements in the lower portion and unsettled fillings below newly provisioned brood cells (Fig.1). For reused nests, we measured the length of the actively used portion (from the lowest newly provisioned brood cell to the nest entrance). Most of our analyses are based on nests in the late phase of the nesting season, between July 15th and August 15th each year).

**Fig. 1:**
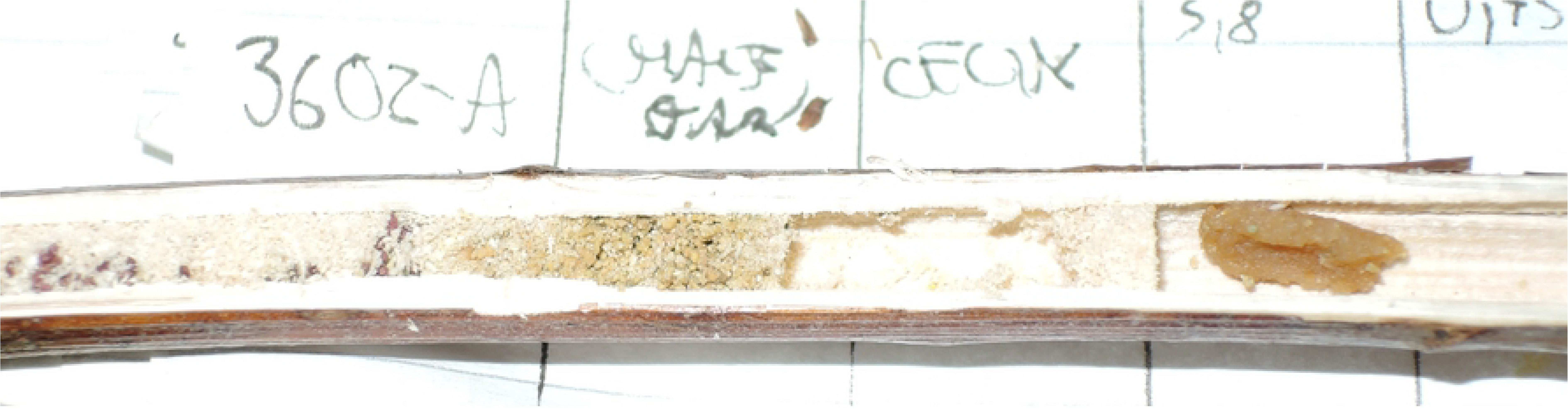
Reused nest of *C. chalybea*. From left to right there are: fillings with excrements of larvae, excrements of young adults, an empty cell, the pollen ball of a brood cell currently being provisioned.

### Classification of nest stage

We only used active brood nests for our analyses. Active brood nests were an outermost brood cell containing a pollen ball or an egg; therefore, all nests that were currently provisioned. We distinguished two types of nests: solitary nests and social nests. Solitary active brood nests contained only a mother and the provisioned brood cells, no young adults. Social nests usually contained a mother and always had at least one young adult in addition to the provisioned brood cells. All together, between July 15th and August 15th we collected 28 social active brood nests (19 in 2015 and 9 in 2017) and 39 solitary active brood nests (19 in 2015 and 20 in 2017). We also classified some nests as full brood nests. Full brood nests were those in which brood cell provisioning had already been completed (the innermost brood cell contained a larva or pupa). Adult offspring are not yet present in these nests, or if they are present, they have not crawled through the nest partitions.

### Observation of social nests

We transported the nests used for observation to special study plots. Transport of these nests was performed in the evening to ensure that all inhabitants were inside the nest. Each study plot contained 24 nets. Here, we only present the results from social *C. chalybea* nests; however, we observed these nests along with nests of other stages and species. Each nest was observed for one observational day, between 8:00 and 16:00 CEST on days with suitable weather. Each plot was observed for the entire time by at least one observer but most of the time there were two observers present. We marked foraging bees with an oil marker (Uni Paint) on the abdomen. We recorded the departure and arrival of foraging bees and noted when adults only departed from a nest (did not return) or newly arrived to a nest. It was necessary to cover the nest entrance with a transparent cup after every arrival of a bee so that the subsequent departure could be observed, since departure is usually very fast. This allowed us to verify whether the bee had been marked previously or if not, to mark it. Nests were dissected after their observational day. We performed this experiment in 2017. We successfully observed 4 social nests of *C. chalybea* per observational day. Two other nests were observed for only a partial day, due to inclement weather conditions.

### Analysis of relatedness between individuals in the nest

We previously developed microsatellites for *C. nigrolabiata* [38], which we also used for the analyses of *C. chalybea*. We used the Chelex protocol for DNA isolation. We isolated DNA from the whole body of eggs and larvae, or the abdomen of pupae and adults. We added 4-8 μl proteinase K and 50 μl of 10% Chelex suspension in ddH_2_0 to dried samples. We mixed the suspension and heated it to 55°C for 45 min, then to 97°C for 8 min in a thermocycler (BioRad). We centrifuged the samples and froze the supernatant for further use.

We used the Type-it Multiplex PCR Master Mix (Quiagen) for multiplex PCR according to the manufacture’s protocol. We used primers for ten microsatellite loci ([38], SI Appendix) at a concentration of 0.05 μmol/l. We used the following settings for PCR: 95°C for 15 minutes; 30 cycles of 94°C for 30 s, 60°C for 90 s, 72°C for 60 s; and finally, 60°C for 30 min.

We mixed 0.8 μl of PCR product with 8.8 μl of formamide and 0.4 μl of marker Liz 500 (Applied Biosystems). We heated the mixture to 95°C for 5 min and let it cool down. Fragmentation analysis was performed on a 16 capillary sequencer at the Laboratory of DNA sequencing at the Faculty of Science, Charles University. We used GeneMarker1.91 software (SoftGenetics, State College, Pennsylvania, USA) for the identification of alleles. Usually, this software correctly identified alleles; however, sometimes manual correction of the size scanner and identification of alleles was necessary.

From the 10 loci we used, 9 provided successful products and all were polymorphic, although polymorphism was highly variable between loci. Loci had between 2 and 33 alleles. Allele frequencies are summarized in [38]. We tested relatedness in 12 social nests. All these nests were collected in 2015. Together, we analyzed 12 old females, 21 non-adult juveniles (15 females and 6 males) and 52 young adults (5 females and 47 males).

### Analysis of relatedness between mother and young adults

In social nests, we used two methods to determine if young individuals in the nest (young adults and also non-adult juveniles) are related to the old female. Primarily, we manually compared the genotypes of the old female and offspring to determine their loci compatibility. Male offspring should contain only alleles that the old female has. Female offspring should share at least one identical allele with the old female at each locus. We counted the number of loci that had alleles that were incompatible with the maternal genotype. We also assessed the relatedness of young adults and offspring in Kinship software [39] using the following analysis: Kinship analysis, relatedness option (Pairwise relatedness: Kinship). As offspring could have a coefficient of relatedness as high as 0.5 and unrelated individuals could have a relatedness coefficient as low as 0, we used 0.25 as the cut-off for related individuals. All offspring below this cut-off were considered unrelated to the old female.

### Testing the maternity of young adult females

To determine the compatibility of young adult females with possible offspring (non-adult juveniles), we counted the number of incompatible loci – loci in the offspring that only had alleles that could not have been inherited from the young adult. We compared the number of loci with incompatible loci between non-adult juveniles and young adult females with the number of incompatible alleles between non-adult juveniles and old females. We also compared the relatedness calculated by Kinship software between non-adult juveniles and young adult females with the number of incompatible loci between non-adult juveniles and old females.

### Testing the paternity of young adult males

We used colony software [40] to test the paternity of young adult males (to the non-adult female offspring) in social nests. Settings in Colony software were: Mating system – Female polygamy, Male polygamy, without inbreeding; Species – Dioecious, Haplodiploid; Length of run – Very long; Analysis method – FL; Likelihood precision – High. For other options, default settings were used. The locus feature was set as all loci codominant. The probability of genotyping error was 0.01; the probability of other errors (for example, mutations) was 0.001 for each locus. The old female was set as the known mother.

### Statistics

All statistical analyses were performed in R software [41]. To test the relationship between sociality and nest reuse, we used Fisher’s exact test. We tested for differences between social and solitary nests with the year as a covariable (model equations were: feature of nest ~ year*sociality). For length of nest, length of nest entrance, and number of brood cells, a linear model was used. For the proportion of empty cells, a generalized linear model of binomial family was used. It was impossible to test sociality and nest reuse in one model together because both factors were strongly correlated. Therefore, we fitted a primary model with sociality (feature of nest ~ year*sociality) and a secondary model with nest reuse (feature of nest ~ year*nest reuse). We compared the Akaike information criteria (AIC) between models with sociality and models with nest reuse for all four tested features of nests. We compared the relatedness between non-adult juveniles and young adult females with the relatedness between non-adult juveniles and old females. We tested differences in the number of incompatible loci by paired Wilcoxon test. We tested differences in relatedness by paired t-test.

## Results

### Evidence for social nesting

In the first part of the nesting season (until July 15^th^), we only observed solitary nests (2013 N=90; 2015 N = 22; 2017 N = 5; 2018 N = 19). None of these nests were reused from the previous season. However, we did find social nests later in the *C. chalybea* nesting season, after the 15^th^ of July. In total, we found 28 social nests. After July 15^th^, half of the active brood nests we found in 2015 were social nests (19/38, Table 1) and 31.03% (9/29) of the active brood nests we found in 2017 were social nests. In 2013 and 2018, all dissected active brood nests were solitary (N=25, 2013; N=26, 2018). One social full brood nest was found in 2018.

**Table 1.** The number of solitary and social active brood nests in different years. Only nests collected between 15 July and 15 August are included.

| Sociality | Solitary nest |  |  | Social nest |  |  |  |
| --- | --- | --- | --- | --- | --- | --- | --- |
| Nest reused? | No | Yes | Together | No | Yes | Together | Total |
| 2013 | 23 | 2 | 25 | 0 | 0 | 0 | 25 |
| 2015 | 15 | 4 | 19 | 3 | 16 | 19 | 38 |
| 2017 | 15 | 5 | 20 | 2 | 7 | 9 | 29 |
| 2018 | 23 | 3 | 26 | 0 | 0 | 0 | 26 |
| Total | 76 | 14 | 90 | 5 | 23 | 28 | 118 |

On average, social nests contained 3.68 young adults (4.15 in 2015 and 2.66 in 2017, Table 2), and the maximum number recorded was 9. Most of the young adults were male (89.32%, 92/103). The reproductive female was present in most nests (82.14%, 23/28), but some social nests had been orphaned.

**Table 2:**
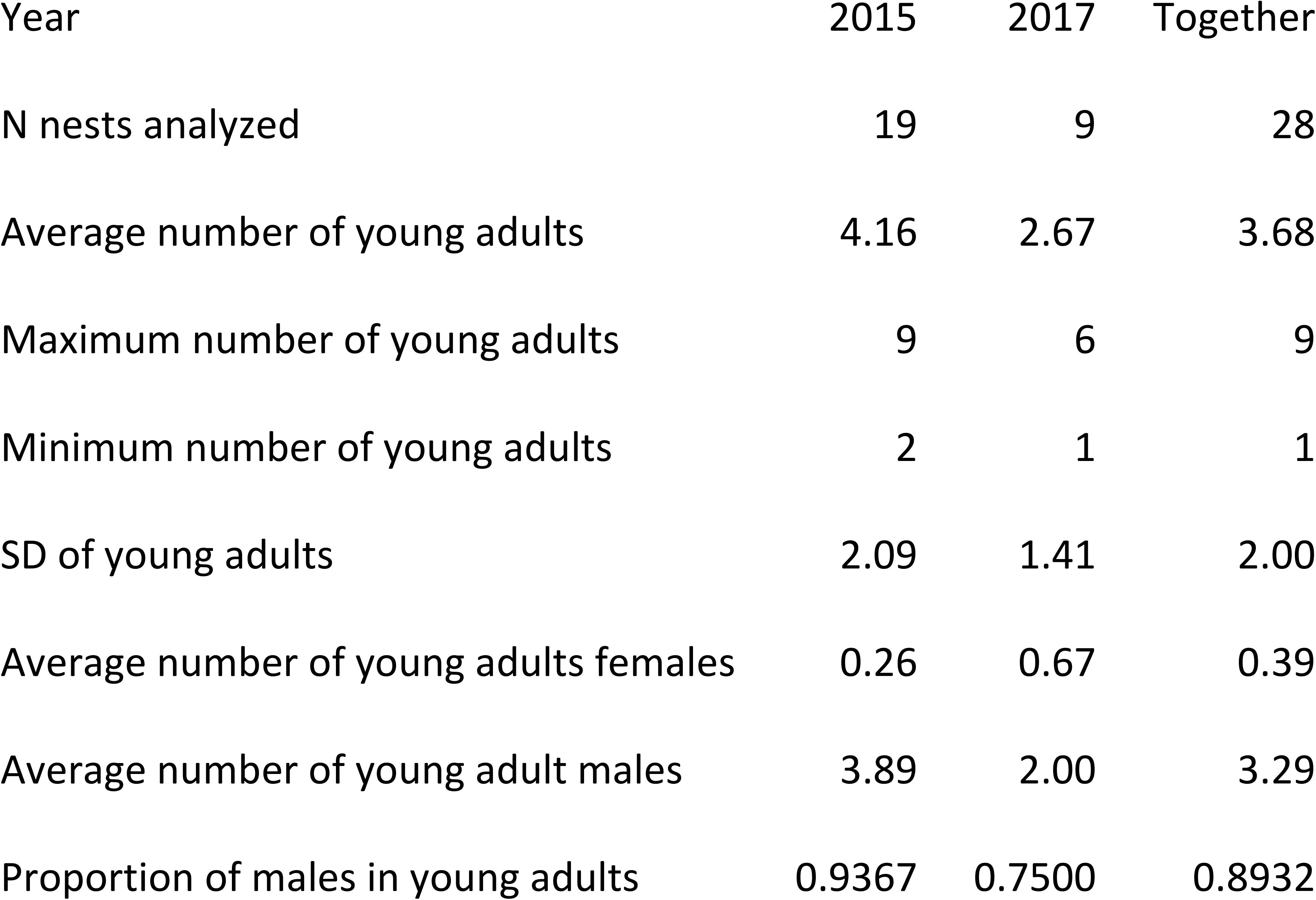
Number of young adults in *C. chalybea* social nests

### Comparison between solitary and social nests

We compared the nest architecture of solitary and social active brood nests. There was a strong association between social nesting and nest reuse: only 15.55% (14/90, Table 1) of solitary nests were reused in comparison to 82.14% (23/28, Table 1) of social nests. This association was significant for both years (Fisher exact test; 2015 – p = 0.0002, N = 38; 2017 – p = 0.0140, N = 29).

Solitary and social active brood nests did not differ in total nest length (linear model, F = 0.81 p = 0.3713, N = 67), but they did significantly differ in the length of the actively used portion of the nest (linear model, F = 17.34, p < 0.0001, N = 67). This result held true when we used nest reuse as the explanatory variable instead of sociality (total nest length did not differ – linear model, F=2.28, p = 0.135, N = 67; but the actively used portion of the nest did significantly differ in length – linear model, F = 43.85, p < 0.0001, N=67). When we compared the AIC of both models, nest reuse was better than sociality as an explanatory variable. There are fillings and excrement from previous instances of nesting at the bottom of reused nests; therefore, the length of the effectively used space is shorter (fig. 1).

We also found a difference in the number of brood cells provisioned. Social nests had significantly fewer provisioned brood cells (linear model, F = 7.21 p = 0.0093, N = 67, Table 3). When we tested nest reuse as an explanatory variable instead of sociality, the difference was also significant (linear model, F = 5.25, p = 0.0253, N = 67); however, the model using sociality had a better AIC than the model with nest reuse.

**Table 3:** Comparison of social and solitary active brood nest characteristics. Only nests collected between 15 July and 15 August are included.

| Nest Type | Solitary active brood nests |  |  | Social active brood nests |  |  | All nests |
| --- | --- | --- | --- | --- | --- | --- | --- |
| Year | 2015 | 2017 | Together | 2015 | 2017 | Together |  |
| Number of Nests Analyzed | 19 | 20 | 39 | 19 | 9 | 28 | 67 |
| Total Length of Nest – mean | 24.53 | 23.32 | 23.91 | 24.31 | 26.83 | 25.12 | 24.41 |
| Length of Nest Used – mean | 22.97 | 20.98 | 21.95 | 14.23 | 17.30 | 15.21 | 19.13 |
| Number of Brood Cells – mean | 3.58 | 2.95 | 3.26 | 2.21 | 2.44 | 2.29 | 2.85 |
| Proportion of Cells Empty – mean | 0.52 | 0.56 | 0.54 | 0.33 | 0.38 | 0.34 | 0.46 |

We also found a difference in the presence of empty cells. In almost all cases, brood cells were separated by empty cells in solitary nests, but they were often adjacent in social nests. The proportion of cells that were empty was significantly lower for social nests compared to solitary nests (Binomial glm, deviance = 7.99, residual deviance = 27.73, p = 0.0047, N = 67, Table 3). When we tested nest reuse as the explanatory variable instead of sociality, there was also a significant difference (Binomial glm, deviance 5.97, residual deviance = 29.78, p = 0.0146, N = 67); however, the model using sociality had a better AIC than the model with nest reuse.

### Foraging activity in social nests

We recorded high foraging activity in social nests, which were observed for a full day (mean = 16.5 foraging trips for a day, range = 12–20, N = 4 nests). In all cases, only ole female performed regular foraging activity. We also observed two additional nests per part of day. In one nest, foraging activity was performed by reproductive female and in second nest no activity was recorded. We did not observe any young adults performing foraging activity; however, we did note the emigration of young adults who did not return to the nest (mean = 1.25 for a day and nest, range = 0–2, N = 4 nests). There was also one case of immigration by a young adult (observed entering the nest without having previously departed).

### Relatedness in social nests

#### Relatedness between the old female and non-adult juveniles (eggs and larvae)

Software analysis in Kinship software concluded that all non-adult juveniles were related to the old female. However, manual comparison of genotypes showed, that one individual had one locus with one allele that could not have been inherited from the old female. Therefore, non-adult juveniles had all loci compatible with the old female’s genotype (95.23%; N = 21, Fig. 2).

**Fig. 2:**
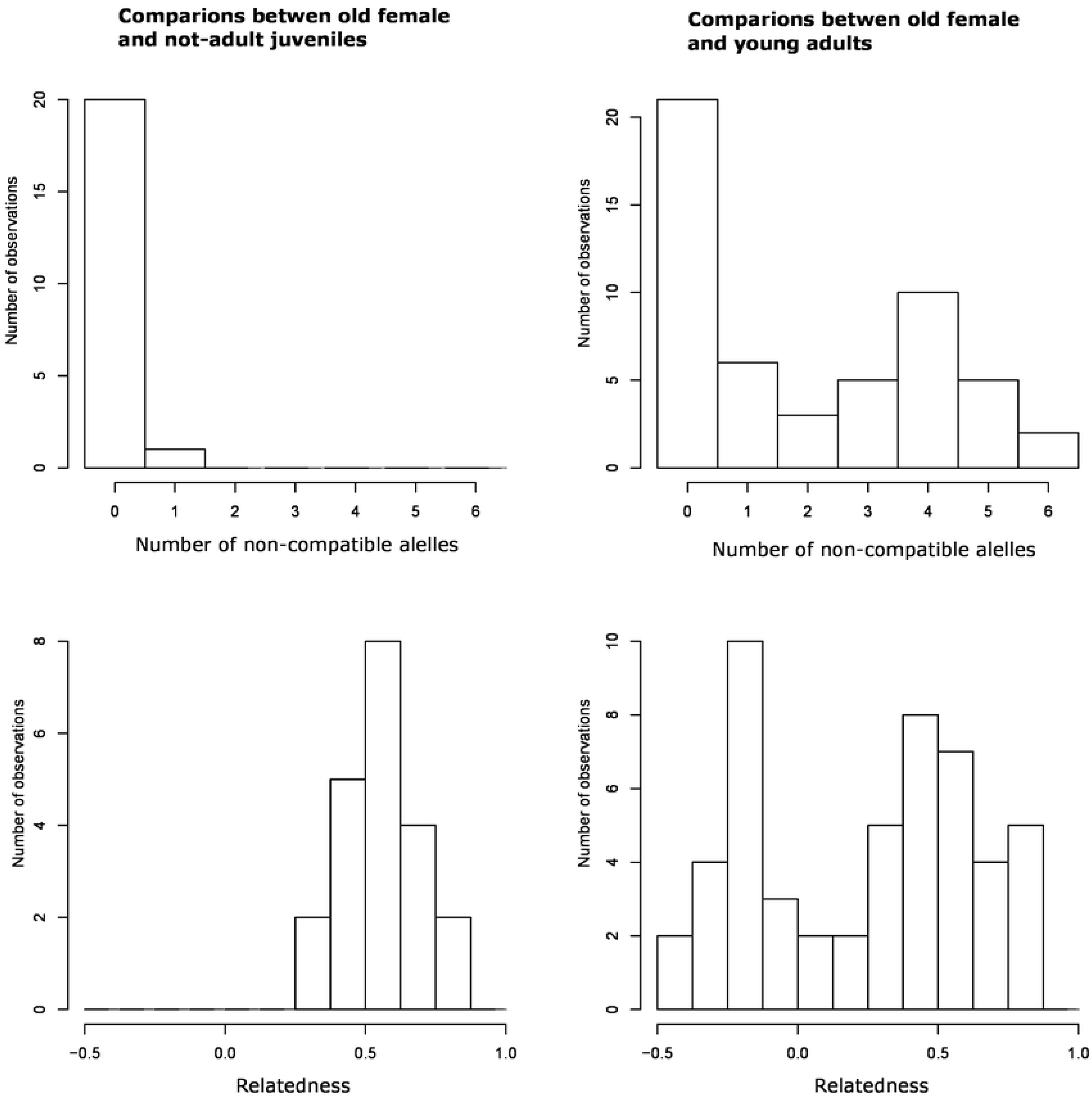
Histograms showing relatedness between the old females and other members of the societies. A) Number of incompatible loci between the old females and non-adult offspring (eggs and larvae), N=21. B) Number of incompatible loci between the old females and young adults, N=52. C) Relatedness between the old females and non-adult offspring (eggs and larvae), calculated using Kingroup software, N=21. D) Relatedness between the old females and young adults, calculated using Kingroup software, N=52.

#### Relatedness between the old females and young adults

Young adults were compatible with the old female’s genotype for all loci in 40.39% of cases (21/52, Fig 2); whereas, one locus disagreed with the old female’s genotype in 11.53% of cases (6/52). For the remaining cases (25/52), the genotyped offspring had more than one locus that disagreed with the old female’s genotype. Analysis using Kinship software showed that 55.77% (29/52, Fig 2) of young adults were related to the old female and 44.23% (23/52, Fig 2) were unrelated to the old female. Therefore, we assume that between 44.23% and 59.61% of young adults are unrelated to the old female.

#### Relatedness between young adult females and non-adult juveniles

Young adult females were only present in 3 out of 12 nests that were genetically analyzed. We compared the relatedness between young adult females and non-adult juveniles with the relatedness between the old female and non-adult juveniles. There were 9 possible pairs of non-adult juveniles and young adult females for evaluation. In 22.22% (2/9) of these possibilities, non-adult juveniles had all loci compatible with maternity of the young adult female; whereas, one locus was incompatible in 33.33% (3/9) of cases and more than one locus was incompatible in 44.44% (4/9) of cases. The genotype of non-adult juvenile offspring was compatible with maternity of the old female in all loci. Therefore, non-adult juveniles have significantly higher compatibility with the genotype of the old female than the young adult females (paired Wilcoxon test, V = 0, p = 0.0206, N=9). Also, we found that the relatedness calculated by Kinship software of non-adult juveniles was higher to old female than young adult females (paired t-test, t = 5.77, df = 8, p = 0.0004, N = 9).

#### Relatedness between young adult males and non-adult juveniles

We tested the paternity of young adult males using Colony software to determine their relatedness to the non-adult juvenile females. This analysis showed that none of these offspring (N = 15) were fathered by any of the young adult males. In all cases, the probability of paternity was less than 1%.

## Discussion

### Sociality in *C. chalybea*

We suggest, that social nests of *C. chalybea* fulfill all three of the conditions for eusociality defined by [24] and [18]: i) reproductive division of labour (only old females reproduce); ii) generation overlap (adults of the parental and filial generations are present); and iii) cooperative brood care (adult members of the colony cooperate in guarding, helping young offspring survive) are all present.

We confirmed the old female’s dominant reproductive role by microsatellite analysis. In all but one case, the genotypes of non-adult juveniles were compatible with maternity of the old female. We suppose that the single locus incompatibility in this one individual is due to a genotyping error or mutation. Young adult females were rarely present in social nests and their maternity is less probable than the maternity of the old female. We also tested that young adult males do not father non-adult juveniles. Therefore, we demonstrated that the old female strongly (probably exclusively) dominates reproduction in social nests and it is highly likely that no other members of the colony reproduce.

The most disputable phenomenon is cooperative brood care. The old female performs all offspring provisioning. We never observed other individuals performing regular foraging activity and we recorded only an occasional emigration or immigration of young adults. We suppose that the presence of young adults is beneficial for nest protection because unprotected nests of *Ceratina* bees [30] and other nest-making social Hymenoptera [42,43] are vulnerable to invasion and destruction.

Social nesting was strongly associated with nest reuse in *C. chalybea*. Nest reuse is generally considered a key factor for the development of sociality in *Ceratina* [36,37,44]. Although nest reuse can be an important factor influencing nest structure in social nests, we showed that sociality itself is a better predictor for the number of brood cells provisioned and the proportion of empty cells in a nest. Therefore, we suggest that at least these aspects of nest use are directly affected by sociality and not only the effects of nest reuse.

### Comparison with other insect societies

The social structure of *C. chalybea* is unusual among social insects for several reasons: i) the presence of young adult males, ii) an unusually high proportion of unrelated colony members, and iii) reproductive subordinate individuals perform only nest guarding (not provisioning).

Almost all (89%) members of *C. chalybea* societies are males. This is interesting because in general, males have a very minor role in the Aculeate Hymenoptera [17,19,20]. A few biparental species are known, in particular crabronid wasps from the genus *Trypoxylon* [45,46] and *Ceratina nigrolabiata* [38]. In almost all eusocial species, males are a small minority among the colony members and have a marginal role in comparison to female workers [21,47]. An interesting exception is the crabronid wasp, *Misrostigmus nigrophtalmus*, in which a high proportion of colony members are male, actively participating in nest defense. They are even able to perform this task in the absence of female helpers [22]. But it remains unknown why male participation in eusocial societies is so rare. Phylogenetic constrains might be one explaination. The solitary ancestors of social species have female care without male participation [17,18,20]. Males lack some morphological structures, such as hairs for pollen collection and a sting, which are important for working effectively in eusocial societies [20,48]. Uncommon male behaviours may arise from performing a task or a standard behaviour with a different primary purpose. In our case, it is likely that males can help with nest protection because they primarily block the nest entrance in self-defense. Regardless of how it occurs, this behavior does lead to effective nest guarding.

We determined that about half of the young adults are unrelated to the old female in *C. chalybea* societies. There exist various mechanisms for arising of insects societies composed of unrelated members, such as pleometrosis [13,15], adopting of orphaned brood [49] or exchange of individuals between neighbour colonies [14]. We can exclude the possibility of pleometrosis for *C. chalybea*, because we never found more than one old female in the nest. Adoption of an unrelated brood is possible because nest usurpation and brood removal do occur in *C. chalybea*; however, it is rare and only occurs with orphaned nests [30]. Therefore, incomplete brood removal cannot explain the large proportion of unrelated young adults in nests of *C. chalybea*. Thus, it is very likely that the unrelated individuals in *C. chalybea* nests originate from neighbouring nests. We frequently observed young adults emigrating from and immigrating to nests; therefore, we consider unrelated individuals to be drifting from other nests.

The reproductively dominant (old) female in C. *chalybea* nests performs all foraging and reproduction. Young adults are passive; they do not perform any regular foraging trips. This type of division of labour is generally uncommon in eusocial Hymenoptera [18,24], but it is usual for Xylocopine bees [25,26,37]. In *C. chalybea*, we found the direct opposite of classical queen-worker task division: the *C. chalybea* old reproductive female performs all foraging trips and young adults (reproductive subordinates) only perform guarding. This is different from Allodapine bees, where multiple females commonly perform some foraging [50], and also from east Asian *Ceratina* of the subgenus *Ceratinidia*, where dominance of reproduction is unstable [51,52]. A direct contradiction of the classical queen-worker roles (foraging dominant, passive subordinate) does occur in *Xylocopa* [26,51,53]. However, in *Xylocopa sulcatipes* societies with a larger number of adult members (about 6), foraging is performed by multiple individuals [53]. Therefore, we have probably found a Hymenopteran society with the lowest proportion of foraging individuals.

### Benefits for young adults

Subordinate members of insect societies usually benefit from indirect fitness [6,9,11]. However, direct fitness benefits, such as the possibility of inheriting a dominant position [25,26,54] or direct reproduction [55,56], can also be important.

Indirect fitness benefits only occur with non-zero relatedness between the donor and acceptor [6]. However, we have shown that about half of the young adults are unrelated to the old female. Moreover, previous work indicates that *C. chalybea* has a multiple mating strategy [38], which is unusual in simple hymenopteran societies [7]. Drifting individuals and multiple mating generate very low relatedness between colony members. Half of the colony members, those that are unrelated, cannot gain any indirect fitness benefit from helping. Furthermore, the other half of the colony members, those that have non-zero relatedness to the colony’s young adults, might only gain a small indirect fitness benefit due to the lower productivity of social nests in comparison to solitary nests.

The possibility for nest inheritance is an important selection factor for the cooperation of unrelated members in small insect societies [54]. Generally, in Xylocopinae bees, nest inheritance is probably a very important driver [25,56]. However, in the case of *C. chalybea* this cannot be an important factor for the retention of sociality, because most of the young adults are male and nest-loyal biparental behaviour is unknown in this species [38]. Additionally, as we did not observe any case of nest reuse from the previous season, therefore we suppose that each female will build new nest next year. For these reasons, we can exclude benefits from nest inheritance as a reason for sociality in the case of *C. chalybea*.

Reproductively subordinate colony members can sometimes directly reproduce in small hymenopteran societies [54,57]. These individuals drifting to foreign nest frequently benefit from direct reproduction and commonly reproduce more than domestic workers [55,58]. However, based on our evidence, it is clear that the old female strongly dominates reproduction in *C. chalybea* societies and reproduction by young adults is negligible or zero. Therefore, direct reproduction cannot be an important motivation for a young adult to stay.

Care of adult offspring is an unusual trait among insects [59], but it is common in Xylocopinae bees [33]. It is likely that young adults remain in their nests because they benefit from the food provided by old female. Long-term cohabitation between an old female and young offspring is a widespread feature in *Ceratina* bees [29]. Many studies of solitary nests have shown that the mother provides pollen and nectar for her young adult offspring [28,31–33]. We argue that social nests arise from nests where mothers feed their adult offspring: first, the mother feeds mature offspring and then she begins to provision new brood cells. However, this strategy can have a significant cost. As provisioning of young adults continues along with providing for new offspring, the mother must divide her resources between the new brood cells and adult offspring; therefore, the amount of food that can be allocated for brood cell provisioning and thus the number of new brood cells is decreased. We observed that the pollen ball in the outermost (open) brood cell, which was currently being provisioned, had an atypical shape in some social nests. We suppose that this pollen ball is partially eaten by young adults. Simultaneous provisioning of brood cells and feeding of young adults has also been documented for *Xylocopa pubescens*. Maternal care of young adults is an important benefit for them [26,60].

Cooperation between organisms is dependent on the costs-benefits ratio [6]. When little cost occurs, little benefit is required to maintain stable cooperation. Young adults of *C. chalybea* do not perform foraging, which is a very risky task for workers in most social insects [18,61]. It is likely that the presence of young adults in *C. chalybea* nests has few costs, because it does not reduce their lifetime reproduction. Females of *C. chalybea* [30] and also other temperate *Ceratina* species do not reproduce before overwintering [29]. Males of temperate *Ceratina* bees survive through the winter and usually mate in the season after overwintering [31,38]. Therefore, remaining in the nest probably has little or no cost to future reproductive success and consequently, only a small amount of benefit is required for young adults to remain.

### Benefits for the old female

There was exceptionally low productivity in social nests of *C. chalybea*. In total, there were fewer brood cells provisioned in social nests than in solitary nests. This differs from other social species, where the overall productivity of social nests is either higher [43,62] or at least the same as solitary nests [44,63]. Contrary to workers in large societies, young adults in *C. chalybea* nests did not leave to perform foraging; rather, they stayed inside their nest. However, non-foraging individuals can be beneficial for the society in other ways. It has been shown that the presence of guards in the nest can be effective protection against pollen robbery by conspecific females in *Xylocopa* [26] or nest usurpation [56].

In comparison to solitary nesting, social nesting decreases the risk of total nest destruction [56,64]. In the case of *C. chalybea*, removal of the mother from completely provisioned solitary nests significantly decreases the survival of offspring due to attack by natural enemies [30]. Therefore, the presence of young adults can be a benefit because they are able to protect the younger cohort of offspring. Young adults can serve the nest community through two mechanisms: i) reducing or eliminating the trade-off between nest guarding and offspring provisioning, and ii) at least temporarily, protecting the nest after the death of the mother. It has been shown that social nesting allows for more effective foraging in multiple facultatively eusocial species [26,65]. We did not test the effectiveness of young adult guarding experimentally; however, we did find a difference between solitary and social nests in their architecture. In social nests, empty cells were significantly less frequent than in solitary nests. Empty cells are thought to be an adaptation for protection against parasite attack [66]. Therefore, in social nests, the presence of young adults can protect against attack and the old female are able to reduce the number of empty cells, allowing more space for provisioned offspring. We observed that young adults were present in some *C. chalybea* nests from which the mothers had already vanished. As these young adults are located in the nest entrance, they can protect the brood cells against potential intruders.

It is possible that there was low nesting productivity in social nests because a significant proportion of pollen and nectar was consumed by the young adults and, therefore, could not be used to build brood cells. As about half of the young adults are related to the old female, feeding of young adults alongside brood provisioning can be beneficial for her reproductive success because this supports their survival [32]. From the old female’s view, social nesting can be interpreted as maternal care for two cohorts of offspring simultaneously: a new cohort of offspring in the brood cells and an old cohort of young adult offspring. However, it is unclear why the old female tolerates unrelated young adults in the nest. One possibility is that it may be difficult to discriminate between alien and own offspring. In *C. calcarata*, the mother can discriminate between nestmate and non-nestmate young females [67]; however, overall aggression among individuals in mature brood nests is generally low, and when it does occur, it is more often against nestmate than non-nestmate young females [67].

### Implications of *C. chalybea* natural history for social evolution

Our observations support the view that benefits for subordinate colony members in small insect societies are not, in many cases, primarily connected to inclusive fitness. It is possible for some females to gain direct fitness benefits, as has been documented in some studies on Xylocopine bees [25,26]. However, in the case of *C. chalybea*, the main benefits are not in the possibility of nest inheritance, but in the extended care of mature offspring. The old female provides pollen and nectar to feed young adults, which helps them survive. The old female tolerates young adults in the nest, because this can provide the benefit of increased nest protection. Therefore, our study supports the importance of mutualistic interactions in the evolution of the early stages of sociality.

As costs to young adults are low, small benefits are sufficient for the maintenance of sociality. We suppose that young adults mainly benefit from the food provided by the old female. Young adults can help with protection against natural enemies; however, their primary motivation for this is probably passive (self-defense). Although the observed society fulfills the definition of eusociality proposed by [18,24],the motivation for the behavior of colony members is mainly selfish. Therefore, the society of *C. chalybea* is something between eusociality and a two-cohort maternal subsociality. Unrelated young adults can be considered parasites, as they take food resources from the old female

Eusociality is ancestral state for all Xylocoinae bees with strict solitarity being a derived strategy [35]. The unusual social organization of *C. chalybea* has some traits in common with typical Xylocopine social organization, especially the presence of unrelated colony members [56,68] and passive reproductively subordinate individuals [25,51]; however, in the quantity of these features, *C. chalybea* is extreme, even among species of the subfamily Xylocopinae. Furthermore, *C. chalybea* society is unique in its inclusion of male colony members.

Here, we have shown that eusociality in bees can be maintained even when the relatedness between colony members is very low and indirect as well as direct fitness benefits (i.e. the possibility of nest inheritance) play small roles. In this case, eusociality is supported by specific natural-history traits (i.e. feeding pollen and nectar to mature offspring and nest reuse). Thus, our results show that good knowledge of natural history is important for interpreting social evolution.

## Acknowledgements

We are grateful to all of the people who helped us with field experiments: Votěch Brož, Kateřina Čermáková, Marcela Dokulilová, Antonín Hlaváček, Lukáš Janošík, Karel Kodejš, Celie Korittová, Eva Matoušková, Zuzana Matějková, Blanka Mikátová, Daniela Reiterová, Jitka Waldhauserová, Šimon Zeman. We are also grateful to Podyjí National Park for permitting our research and for their friendliness. We also thank the Catholic Priest, Marian Hušek, for accommodation during our field research in the rectory in Havraníky village.

The Grant Agency of Charles University in Prague (Grant GAUK 764119/2019) and the Specific University Research project Integrative Animal Biology (Grant SVV 260434/2018) supported this research.

## Data Availability

Dataset is available as SI material of this paper.

## Author contribution

MM and JS designed the research; MM, DB and JS performed the research; MM analyzed the data; MM wrote the initial draft of the paper; all autors commented and finalized the paper.

